# Case report on lethal dog attacks on adult rhesus macaques (*Macaca mulatta*) in an anthropogenic landscape

**DOI:** 10.1101/2023.08.16.553455

**Authors:** Bidisha Chakraborty, Krishna Pithva, Subham Mohanty, Brenda McCowan

## Abstract

For nonhuman primates living in anthropogenic areas, predation by larger predators is relatively rare. However, smaller predators such as free-ranging as well as domesticated dogs can shape the socioecology of urban nonhuman primates, directly by attacking and predating upon them, or indirectly by modifying their activity patterns. Here, we describe 3 (2 potentially lethal) cases of dog attacks on adult rhesus macaques inhabiting an anthropogenic landscape in Northern India, and the circumstances surrounding these incidents. We discuss the importance of considering the presence of dogs while studying nonhuman primate populations across the anthropogenic gradient and its implications for understanding how human presence can directly and indirectly affect predator-prey relationship in these areas, as well as its potential role in modifying group social dynamics as well as in transmission of zoonotic agents.

## INTRODUCTION

In response to human population-induced habitat alteration, nonhuman primate species are either getting locally extinct (Jones-Engel et al., 2022) or adapting themselves by being more ecologically flexible while being forced to live in close proximity to humans (Hockings et al., 2012). Animals living in anthropogenic areas need to balance novel costs and benefits associated with such areas to survive. Living among humans comes with costs, such as elevated stress levels (Maréchal et al., 2011), increased aggression from conspecifics (Southwick et al., 1976) and humans (McCarthy et al., 2009), human intervention in the form of culling, translocation (Massei et al., 2010), death from electrocution (Pereira et al., 2020) and roadkills (Hetman et al., 2019). Though the largely predictable nature of high-quality anthropogenic food can positively affect individual fitness (Kurita et al., 2008), anthropogenic food like garbage and junk food can also negatively affect animal cardiovascular health (Hannah et al., 1991).

A potential benefit of living in anthropogenic areas is thought to be reduced predation risk due to decreased presence of large predators such as tigers (*Panthera tigris*) and leopards (*Panthera purdus*) (Crooks & Soulé, 1999). Lack of such top-down competition, in turn, positively affects the population of mesopredators such as coyotes (*Canis latrans*), dogs (*Canis lupus familiaris*), racoons (*Procyon lotor*) and red foxes (*Vulpes vulpes*) (Prugh et al., 2009), which might affect certain prey species (Takimoto & Nishijima, 2022). Being one of most abundant carnivores across the globe, dogs pose significant threats to wildlife conservation by disturbing, competing with and predating upon vulnerable wildlife (Gompper, 2013) and can directly affect predator-prey dynamics in anthropogenic areas (Dorresteijn et al., 2015). Human-dog coexistence has been observed for a long time, and dogs have been used by humans for protection of crops, watching over livestock and in hunting (McKinney et al., 2023; Miller et al., 2013). Domestication of dogs, as well as the tolerated presence of free-ranging dogs can not only directly affect local wildlife by predation but can also have non-consumptive effects by affecting food and space utilization (Gompper, 2014) and altering time-activity budgets (Waters et al., 2017). Moreover, since dogs interact with both humans and wildlife, they can also act as potential zoonotic agents, aiding in the bidirectional transmission of several bacterial (Chomel, 2014), protozoan (Oliveira et al., 2019) and viral (Sun et al., 2010) diseases to humans and other wildlife. In fact, incidences of canine rabies in humans after nonhuman primate-related injuries have been reported across the world (Gautret et al., 2014; Kumar Bharti, 2016) implying occurrences of cross-species spillover. In some Asian countries, high incidence of rabies in rhesus macaques *(Macaca mulatta)* can be attributed to coexistence with stray dogs, also making humans susceptible to the virus through bites and scratches from both species (Taku et al., 2007).

Rhesus macaques *(Macaca mulatta)* have been one of the most successful nonhuman species to adapt to anthropogenic pressures and coexist to some extent in human-impacted landscapes, occupying broad ecological niches ranging across forests to cities. Given their socioecological flexibility, their population has been steadily increasing in urban landscapes, often resulting in competition with humans for resources such as food and space (Cooper et al., 2022). Larger predators of rhesus macaques like tigers and leopards usually avoid dense urban areas (Carter et al., 2015), while cooccurrences of dogs and rhesus macaques, often leading to aggression, is often observed, and depredation of rhesus macaque infants by dogs has also been reported (Anderson, 1986).

Cases of predation or harassment by dogs on nonhuman primates have been reported across the primate order. Predation and attacks on brown howler monkey (*Alouatta guariba*) (da Silva et al., 2021), black howler monkeys (*Aloutta pigra*) (Franquesa-Soler et al., 2023), Japanese macaques (*Macaca fuscata*) infants (Hill, 2014), young female and infant barbary macaques (*Macaca sylvanus*) (Waters et al., 2017), juvenile long tailed macaques (*Macaca fascicularis*) (Riley et al., 2015) and Central Himalayan langurs (*Semnopithecus schistaceus*) (Nautiyal et al., 2023) have been recently reported, along with several other cases from earlier (reviewed in Anderson, 1986). Despite both being some of the most widespread non-human animals, surprisingly little is known (or reported) about rhesus macaque-dog interactions. Given the role of dogs as predators of nonhuman primates, detailed records of their interactions are needed to understand their role (if any) in shaping the socioecology of nonhuman primates in human-impacted landscapes. In this report, we add to the growing literature on dog-non-human primate interactions by sharing 3 cases of dog attacks (2 of them potentially being fatal) on rhesus macaque adults in an anthropogenic landscape in North India. To our knowledge, this is the first reported case of fatal dog attack on rhesus macaque adult(s).

## STUDY SITE AND SUBJECTS

This study was set in a Hindu temple called Jakhu Temple in the city of Shimla in Himachal Pradesh in North India (31.1008° N, 77.1845° E). This site (approximately 0.02 km^2^) is frequented by around 5-6 groups of rhesus macaques as well as hundreds of tourists every day. The site consists of paved temple premises, a central garden area, a few small restaurants, a cable car tower, and a guesthouse attached to the temple grounds. Descending from the main temple are forested slopes on all sides as well as stairs on one of those slopes leading down to more densely populated areas (for a more detailed description of this site, see Kaburu et al., 2019). The site is also home to 4-5 semi-domesticated or owned free-roaming dogs that are seen across the site. This population was first monitored from 2016-2018 as part of a project exploring human-macaque interactions and macaque social behavior (Kaburu et al., 2018) where 3 groups of rhesus macaques were studied. Following that, one of the original study groups (Shaggy’s Group or SG), consisting of 48 adult individuals (38 females and 10 males) was followed from April 2022-June 2023 on a project exploring intergroup conflict, for which detailed behavioral and feeding data is being collected.

## BEHAVIORAL DATA COLLECTION

The study group (SG) was followed from 9 AM-4:30 PM 4-5 days a week recording detailed individual and group-level data on social behavior, feeding behavior as well as human-macaque interactions. We also recorded the presence/absence of dogs during our data collection. The three cases of dog attacks on rhesus macaques reported here were opportunistically recorded using video recording aided with narration.

## RESULTS

### Overall nature of the dog-rhesus macaque interactions

Dogs were present at this site during the observation period in 2016-2018, but the nature of dog-rhesus macaque interactions overall seems to have become more aggressive. 4 large and one small dog was commonly observed at the site during the current data collection period (2022-23), with 3 of the dogs belonging to a local family, with the other was free-ranging. When one or more of the larger dogs were present, the macaques were seen running away, trying to get to higher ground (trees or temple walls). Some macaques would threaten the dogs from a distance, some even attempting to slap or scratch the dogs. After interactions with dogs, the macaques would stay on higher ground for a while. Interestingly, this behavior would change if the dog was smaller. Usually, the macaques would ignore the smaller dog, and even chase it. Differential behavior based on the dogs’ characteristics has also been reported in vervet monkeys (*Cercopithecus aethiops*) (Galat & Galat-Luong, 1977). Often, the dogs were also used by local watchmen to chase monkeys away from tourists or to break up macaque fights. Here we present the detailed descriptions of the 3 cases of dog attacks on adult rhesus macaques. For summary of circumstances surrounding each incident along with the behavior of the attacked individual and their conspecifics, refer to Table 1 of SI.

### Case-1: 31st August, 2022

An adult female from SG (Daisy) was observed with a small head injury in May 2022, which expanded in size and progressively got worse over time, eventually rendering her unable to walk normally (Figure 1a). On 31st August 2022 at around 4:15 PM, we observed a fatal dog attack on this female. The event took place in the parking lot of the study site, where 2 other groups were present (PG and WG) at the time. During the event, 3 dogs were chasing the monkeys, which caused them to climb up trees on the nearby slopes. Individuals from our study group (SG) were dispersed, with some being on trees on the slope, some on the roof of another nearby temple looking down at the parking lot, and some being in the main temple premises away from the parking lot. While attempting to escape the dogs, the female (Daisy) fell from a slope onto a car from a height of 8-10 meters, following which she tried to escape to the nearby slope. However, given her slow gait, she was captured by one of the dogs that bit and dragged her down to the slope. There were around 20 individuals from SG (15 females and 5 males) present at the scene, with some watching the event and emitting alarm calls from the nearby temple roof, while 5 individuals (4 mid-high ranking adult males from SG and 1 mid-ranking adult female) went closer to where it was occurring and started emitting loud alarm calls. Individuals from one of the other groups (WG) fled to higher ground and watched the incident from the trees, while individuals from another group (PG), specifically 1 adult male and 2 adult females went close to where Daisy and the other dogs were and started emitting loud alarm calls. After 10-12 minutes, when individuals from all 3 groups (SG, PG and WG) left, we looked for the body that was down the slope. The female seemed to be barely alive, with slight twitching of hands, and flies settling on her head and limbs. After a while, it started raining and we could not see her moving anymore. While the dog attack was taking place at the parking lot, around 9 individuals (3 males and 6 females) that were observed at the main temple away from the event were seen looking at the general direction of the rest of the group and making contact calls.

**Figure 1a:**
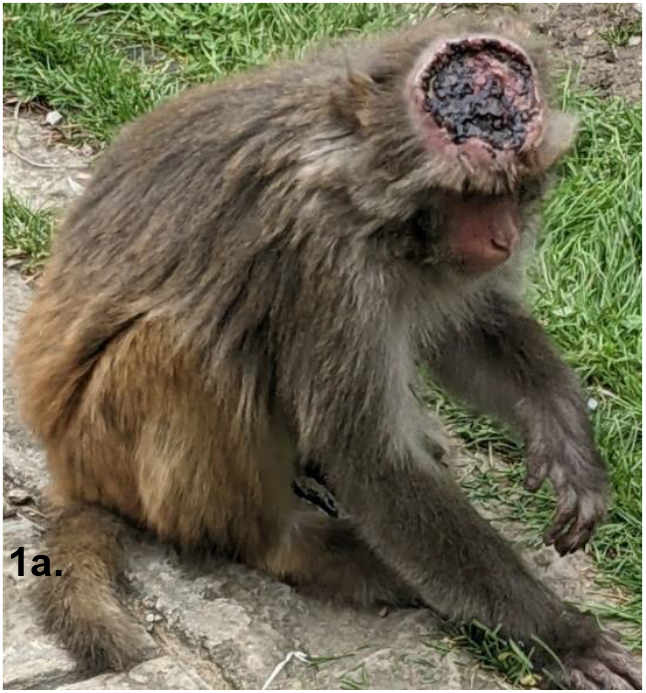
Daisy’s injury.

The next day, 1st September 2022, the corpse was still on the same slope (Figure 1b). Some individuals from SG went down to the road adjacent to the body. The second highest ranking male in SG sat for about 10 minutes on the railing on the side of the road looking down the slope. After he left, individuals from the other group (PG) spent 10-15 minutes looking down the slope.

**Figure 1b:**
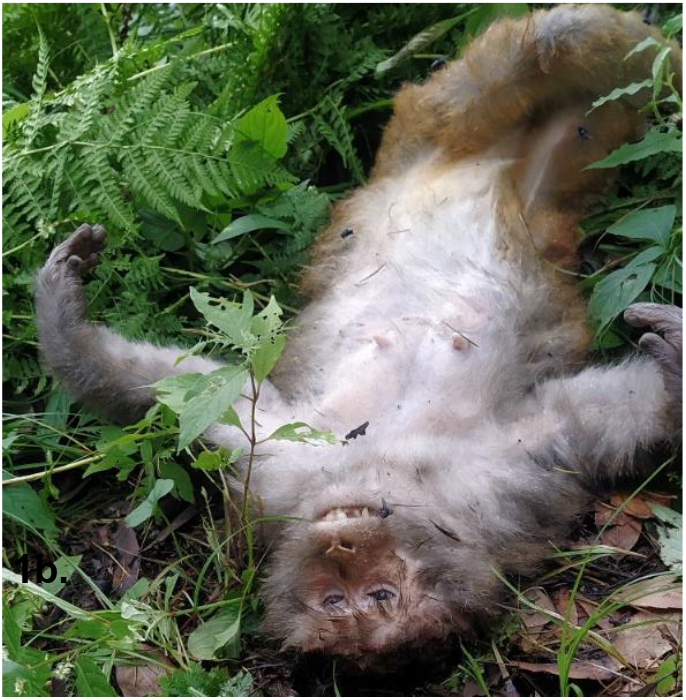
The corpse of the deceased female.

### Case-2: 28^th^ November, 2022

Around 12:05 PM on 28th November, an intergroup conflict (IGC) started between our focal group (SG) and another group (GG). While the IGC was taking place, 4 dogs (3 large and one small) attacked and bit a female from GG (video provided in *SI*). Initially, around 5-6 individuals from our study group (SG) were present along with around 6 individuals from GG. As soon as the female was bitten, both groups ran towards the scene and climbed up trees while emitting vocal threats towards the dogs and their conspecifics, as well as alarms calls. One of the larger dogs kept biting the female and had the female immobilized by the neck for 20-30 seconds (Figure 2a). If any monkey tried to go close to the female, they were chased away by the other dogs. This event went on for 4-5 mins before a local watchman was able to chase the dogs away. Both groups kept emitting alarm calls from higher ground throughout. The female had a gaping, bleeding wound across her thigh to her abdomen, as well as wounds on her left knee, shoulders, neck, right feet and anogenital area (Figure 2b). 5-6 males remained close to her and aggressively threatened anyone nearby. The female kept licking her injury and eventually dragged herself down the closest slope since she could not walk. 2 females from the group tried to get close to her, and one of them tried to inspect her wounds and groom her, but the female slowly dragged herself deeper into the forested slope where we lost sight of her. During this time, males from our study group aggressively lunged and chased the dogs away. Following this event, the female’s group (GG) were seen quietly going down the slope, while monkeys from our study group (SG) were uncharacteristically silent while they rested and groomed on higher ground.

**Figure 2a.**
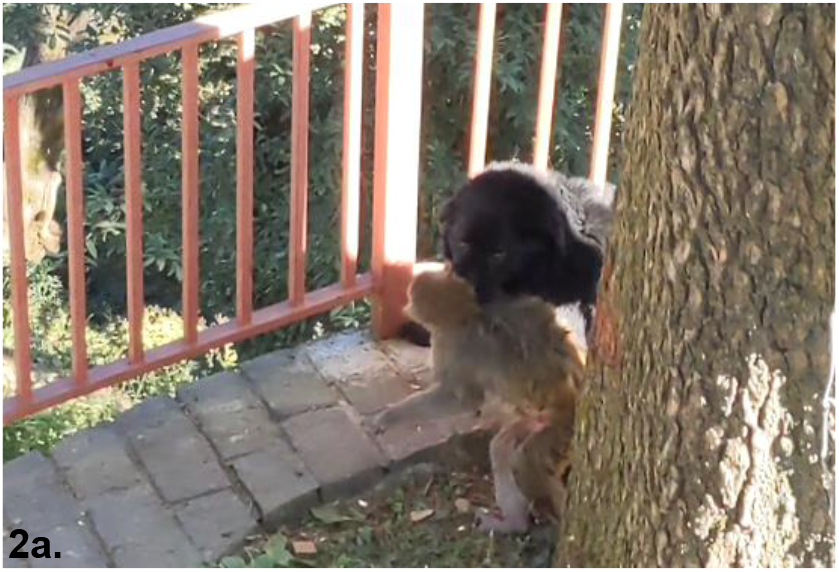
Injuries from the dog attack.

**Figure 2b.**
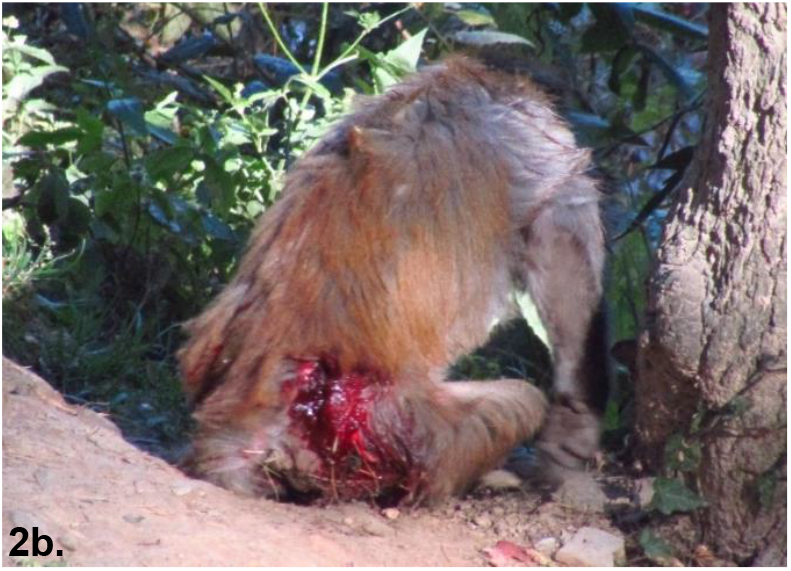
The dog biting the female’s neck.

The next day, we did not observe GG near the site, but a few individuals (1 male and 2 females) from our study group went down the slope where the female was last seen, and started emitting alarm calls. Later, we tried to go down the slope to potentially locate her corpse, but given the dense vegetation, it was not possible to do so. Moreover, since animals’ conspecifics can aggressively defend their corpses (Buhl et al., 2012), we decided that it would not be safe to venture closer.

The female was not seen with GG after that day. Sometimes sick and injured individuals distance themselves from their conspecifics (Stockmaier et al., 2021), so she might have stayed at the periphery of the group. However, despite us monitoring the group for months following the event, no female with such extensive injuries was observed again, suggesting a high likelihood that the female succumbed to her injuries.

### Case-3: 8^*th*^ December 2022

Around 10 AM on 8th December, we witnessed an intergroup conflict (IGC) event between our study group (SG) and another group (RG) at the site. The fight started at the main temple area and the groups eventually chased each other down the neighboring slopes. Initially, 13 adults and 3 juveniles were present from SG, while 13 adults and 8 juveniles were seen from RG. At around 3-4 mins after the IGC started, we heard alarm calls from both the groups, following which 3 dogs ran to the area and chased the monkeys down the slopes. A male from the RG was attacked by a dog as he tried to descend from a tree and run down the slope. The dog was seen to momentarily bite the male’s left abdominal area, but the male was able to break free and escaped. Most of the individuals present from both groups ran up trees on the slope and emitted alarm calls for a while. We were not able to identify the male since he swiftly escaped and hence, were not able to observe this individual’s behavior later.

## DISCUSSION

In this paper, we report 3 cases of dog attacks on adult rhesus macaques at an anthropogenic site in India, and the first reported case of fatal dog attack on r. In two of these cases, two adult females were seriously injured and one (potentially both of them) died as a result of the attack. Albeit limited, these observations and differences in outcome of the attacks can highlight the differential tradeoffs faced by individuals of varying sociodemographic classes. For females, life in anthropogenic areas might be especially costly given their life-history traits. They are generally smaller in size and often carry dependent offspring, thus making them especially vulnerable to dog attacks. Similar instances of smaller and more vulnerable age-sex classes falling prey to dogs have been reported in other cases too (long-tailed macaques: Riley et al., 2015; barbary macaques: Waters et al., 2017).

Apart from lethal consequences of interaction with dogs, the mere presence of dogs can substantially impact the socioecology of nonhuman primates. Presence of dogs negatively affects social behavior and foraging activities by imposing time and opportunity costs (Gumert et al., 2013; Riley et al., 2015). Direct interactions with dogs as well as anti-predator vigilance activities can cause physiological stress (Rangel-Negrín et al., 2023) and eventually might even have fitness effects. In fact, presence of dogs seemed to negatively affect juvenile population sizes in a population of long-tailed macaques (Gumert et al., 2013). Due to these factors, dogs have been referred to as serious predators for many urban/peri-urban long-tailed macaque populations (Riley et al., 2015). One of the socioecological models (series of theoretical models predicting variation in grouping patterns in primates and other mammalian species) predict that predation risk is a major factor driving intergroup and intragroup behavior (Schaik, 1989). Despite the widespread presence of dogs alongside nonhuman primate populations and their effect on nonhuman primate behavior, the effect of smaller predators like dogs are rarely examined in the context of nonhuman primate socioecology. Whether dogs in anthropogenic areas impart as much selective pressure as predators in more wild populations is yet to be tested and can only be elucidated by accounting for their presence in such populations while studying their socioecology.

Observations such as ours can also be used to speculate on the circumstances surrounding dog attacks and their implications for the management of dog-human-nonhuman primate interactions at these anthropogenic interfaces. Anthropogenic areas often have clumped food sources, such as areas of provisioning and garbage, where humans and animals frequently aggregate, often leading to aggressive interactions. All three of the dog attacks reported here were preceded by intergroup interactions between two or more groups. Given the fast-paced nature of these interactions, it is hard to say if the dogs responded to the sounds of the macaques or if they responded to the sounds given by temple guards in an attempt to break up the intergroup fight or a combination of both. Dog-monkey interactions during foraging events in anthropogenic areas have been reported in vervet monkeys (Butler et al., 2004) as well long-tailed macaques (Riley et al., 2015). Given the frequent tridirectional interactions of synanthropic nonhuman primates with humans as well as dogs, pinpointing spatiotemporal factors driving these multispecies contact patterns might help us pinpoint hotspots of such encounters and manage the potentially harmful consequences of such interactions like physical harm caused to both species and bidirectional zoonotic transfer.

Such reports can also offer us insights into how individual and group behavior is manifested during such relatively rare incidents. We observed differences in the behavior of the group members to the death of the two females. In the case of the first female from our study group (Daisy) who had been sick for a while, her corpse received very little interest from her conspecifics. In contrast, the second female’s traumatic death elicited aggressive displays from her conspecifics. Such difference in the behavior of conspecifics and mothers based on the ‘peaceful’ versus traumatic death has been reported in many nonhuman primates such as rhesus macaques (Buhl et al., 2012), Japanese macaque (Sugiyama et al., 2009), chimpanzees (*Pan troglodytes*) (Cronin et al., 2011) and others (Anderson, 2011) and is believed to be influenced by the tangible nature of traumatic deaths (usually infanticide or predation) which conspecifics can physically witness and intervene in, versus sickness (Anderson, 2011). Moreover, death of group members can affect intra-group affiliative (Buhl et al., 2012) and aggressive behavior (Kaburu et al., 2013), but this can be more pronounced if the deceased individual occupied a socially central or dominant position in the group (Kaburu et al., 2013). The deceased female from our study group was low-ranking and socially peripheral, and we did not observe any obvious group-level changes. However, future analysis will explore intragroup behavioral changes after her death, as well as changes in social dynamics of her affiliative partners.

Overall, this report will add to the growing literature on how urban predators such as dogs can impose significant effects on the survival and behavior of synanthropic nonhuman primates, and thus are important entities to consider while studying (peri)urban nonhuman primate populations. Moreover, given the important implications of these interactions for zoonotic disease transmission, such reports are vital in our understanding of complex pathways of potential biological agents.

## ACKNOWLEDGEMENTS

We thank the Himachal Pradesh Forest Department for their permission to conduct research at this site. We also thank Dr. Stefano S.K. Kaburu and Dr. Michelle Brown for their constant support and advice regarding data collection. We are also grateful to the Leakey Foundation, Primate Society of Great Britain, UC Davis Richard Coss Award for their generous financial support. We also thank Dr. Sandeep K. Rattan for his help and ongoing interest in this research. Finally, we would like to express our gratitude to members of the McCowan lab at UC Davis for their helpful advice and suggestions regarding this manuscript.

## COMPLIANCE WITH ETHICAL STANDARDS

This study was completely observational and did not involve interaction or experiments with the study animals. This study was also conducted in collaboration and consultation with the Himachal Pradesh Forest Department and adhered to the local Indian laws. The authors do not have any conflict of interest.

## Notes

### Competing Interest Statement

The authors have declared no competing interest.

