## supplemental table for "Case report on lethal dog attacks on adult rhesus macaques (*Macaca mulatta*) in an anthropogenic landscape"

Bidisha Chakraborty^1^, Krishna Pithva^1^, Subham Mohanty^1^, Brenda McCowan^1,2
1^Department of Population Health & Reproduction, School of Veterinary Medicine, University of California, Davis, CA, USA
^2^California National Primate Research Center, University of California, Davis, CA, USA

Correspondence: Bidisha Chakraborty (ORCID ID: 0000-0003-2458-0647)
Department of Population Health & Reproduction, School of Veterinary Medicine, University of California, Davis, CA, USA,

**Table 1**: Summary of the 3 dog attack events

| **Event** | **Attacked individual** | **Context of incident** | **No. of dogs** | **Monkey to dog behavior** | **Dog to monkey behavior** | **Monkey to monkey behavior** | **Human intervention to the attack** | **Outcome** |
| --- | --- | --- | --- | --- | --- | --- | --- | --- |
| 1 | An adult female (Daisy) from study group (SG) | Intergroup aggregation of study group (SG) and 2 other groups | 3 | Ran away, climbed up trees, emitted alarm calls. | Dogs chased monkeys down the slopes. An adult female was bitten by a dog. | Contact calls emitted by individuals from the attacked female's group. Individuals from the female's as well as other groups visited the attack site. Contact calls emitted by group conspecifics. | None | Presumed internal injuries due to falling from the parking lot. Bitten by dog. Later found dead. |
| 2 (video provided) | An adult female from non-study group (GG) | During intergroup conflict | 4 | Ras away, climbed up trees, emitted alarm calls, aggressive vocalization towards dogs. Some individuals chased dogs. | Dogs chased monkeys away. An adult female was bitten and attacked by 4 dogs. | Aggressive vocalizations directed towards conspecifics from their own and other groups. Males from both groups threatened everybody in the proximity. Individuals from the female's group attempted to approach and groom the attacked female. | Local watchman intervened by chasing dogs and monkeys away using sticks and by bursting crackers | Severe wounding and gaping, bleeding across the thigh to her abdomen. Wounding on her left knee, shoulder, neck, right feet and anogenital area. Presumed dead |
| 3 | A male from non-study group (RG) | During intergroup conflict | 3 | Running away, climbing trees, emitting alarm calls. | Dogs chased monkeys, one dog bit an adult male. | Individuals from both groups stayed on trees and emitted alarm calls for a while. | None | Bitten on the left side of abdominal area but no visible injuries seen. |
